# Human Rhinoviruses: a novel class of oncolytic virus

**DOI:** 10.1101/2023.05.31.542867

**Authors:** William J Burnett, Amber Cluff, Melissa Reeves, Gennie L Parkman, Chase Hart, Andrew Ramstead, Sheri Holmen, Matthew A Williams, Matthew W VanBrocklin

## Abstract

While continuing to develop new and improved OVs, researchers have been vigilant in optimizing their ability to safely infect, replicate, and induce an immune response within tumors. There are currently two crucial limitations in OV development, however: modification of viruses often negatively impact their ability to infect, and induction of strong immune responses also promotes rapid viral clearance. In this study, we investigate wild-type HRVs as a novel class of OV. Multiple HRV serotypes were propagated in human melanoma cell lines to produce highly oncolytic populations of virus. A large panel of cancer types were infected, with cytotoxicity evaluated using flow cytometry and real time live imaging. Pro-inflammatory signaling was assessed by cytokine multiplexing. Tumor responses to HRV were assessed in human xenograft and in syngeneic, immune-competent mouse tumor models. We find that HRVs are capable of infecting and killing a wide variety of human cancer cell lines *in vitro* and *in vivo*, inducing pro-inflammatory responses and, ultimately, tumor regression. We propose that the natural safety profile of these viruses, coupled with their anti-tumor efficacy and multivalent potential seen in our preclinical model systems, make HRVs ideal candidates for development as oncolytic viruses for clinical testing.

**Simple Summary:** The idea of utilizing viruses to combat cancer has been around for over a century, yet in practice has seen relatively modest therapeutic success following decades of strategic design to improve safety, efficacy, and tumor selectivity. Engagement of an immune response is critical to have lasting tumor regression upon treatment, but many patients are unable to mount a sufficient response to standard-of-care immunotherapy alone. Oncolytic viruses (OVs) have been shown to synergize with such therapies to accomplish this goal to varying degrees. The purpose of this study is to investigate human rhinoviruses (HRVs) to establish their efficacy as a novel class of anti-cancer agents. We aim to use HRVs to promote longer lasting and more effective immune responses to tumors by fostering viral persistence and immune evasion, in addition to developing multivalent treatment strategies that enhance tumor regression by preventing tumor escape from viral infection.

## Introduction

At the beginning of the twentieth century, by Dr. George Dock observed something peculiar: leukemia regression in a patient apparently infected by a strain of influenza virus [1-3]. However, since the field of virology was barely beginning to emerge at this time, follow-up studies were hindered. A strong resurgence of interest in oncolytic viruses would occur decades later when cell culture and *ex vivo* viral propagation became common [2]. In the 1990’s, major advances in sequencing and recombineering technologies would further advance the field by allowing the generation of attenuated and modified oncolytic viruses, such as herpes simplex virus-1 (HSV-1), that preferentially infect tumor cells [1, 4]. While some viruses were approved for use in human cancers by other regulatory agencies previously, Talimogene laherparepvec (T-VEC) was the first oncolytic virus approved by the Food and Drug Administration in the United States in 2015 [5]. Incredibly, there are now several families of viruses, encompassing dozens of variations, that have successfully transitioned into early-phase clinical trials [1]. In the United States alone, there are over 100 ongoing clinical trials evaluating single agent oncolytic viruses or viruses in combination with other treatment modalities in many cancer types.

With some exceptions, OVs have not been overwhelmingly impressive as single agents in early phase clinical trials. Nevertheless, the advent of immune checkpoint blockade (ICB) immunotherapies, along with their clinical success in melanoma and other cancers, has given new life to OV research. Importantly, ICB and OVs have non-overlapping functions. Treatment with a checkpoint inhibitor results in the disinhibition of CD8+ T-cells, which drives tumor recognition and targeted elimination [6]. Infection with a virus kills and lyses tumor cells, driving an antigenic form of cell death that promotes upregulation of interferon response genes [6]. This consequently enhances recruitment of immune cells to the site of infection, increase tumor antigen presentation by antigen presenting cells (APCs), and upregulate PD-L1 on the surface of the tumor. When combined, these treatment strategies have demonstrated potent synergy [6-8].

While developing new and improved OVs, researchers have been vigilant in optimizing their ability to infect, replicate, and induce an immune response, while simultaneously striving to minimize toxicity. As an illustration of this, T-VEC is genetically modified to enhance its properties as an OV. ICP34.5 is deleted to mitigate toxicity by impairing neurovirulance and pathogenicity [9-12]. Additional modifications have been made to enhance the immune response to the viral infection, including deletion of ICP47, which acts to suppress antigen presentation. This suppression of antigen presentation is a key factor in mounting adaptive immune responses [13, 14]. Deletion of ICP47, therefore, results in increased tumor cell recognition by CD8+ T-cells. T-VEC has also been armed with human granulocyte macrophage colony stimulating factor (GM-CSF) with the aim of improving tumor antigen presentation by professional APCs [15]. However, despite its FDA-approved status, T-VEC maintains limited application. As a herpes-simplex virus (HSV-1) that infects neurons, T-VEC cannot be delivered systemically and thus is restricted to injectable lesions. The alterations described are designed to restrict its replication to tumor cells, but the cost of attenuation is to limit its natural ability to effectively infect and propagate, even within tumors. Furthermore, as with other OVs, induction of strong immune responses also promotes rapid viral clearance, which may diminish its anti-tumor effect [1, 16].

It would be beneficial to identify naturally occurring viruses that would not require further modifications to be efficacious and safe. In this way, the virus’ natural replicative and infective properties could be retained. Desirable features for oncolytic viruses include an inherent tumor tropism, rapid replication within tumor cells, induction of immunogenic cell death, and viral persistence with low toxicity.

Human rhinoviruses (HRVs) are members of the picornavirus family and hold the title of the most common infectious agent known humankind [17]. They are the major cause of the “common cold” via infection of the upper respiratory tract and are estimated to be responsible for up to half of all human infections [17]. Despite this impressive pathogenicity, in a study of short-term repeated HRV exposure, most (82%), but not all, human subjects infected with high titers of HRV caught colds, even when exposed directly through the upper respiratory tract [18]. Symptoms from infection tend to be mild and HRVs are eventually cleared from the body, though it has proven incredibly challenging, if not impossible, to generate effective vaccines [19-21]. This is due to the dynamic evolution of HRV, which occurs via genomic recombination and low fidelity replication (6.9×10^−5^ substitutions/nucleotide/cell infection), generating an incredible amount of diversity [21-24]. While neutralizing antibodies to HRVs are detected upon infection, antibody titers rapidly decrease over time and are not retained in the body long-term. This contrasts with what is seen during infection by other picornaviruses, such as coxsackievirus and poliovirus [25]. There is also little cross-reactivity between antibodies against one serotype of HRV versus another [19]. Taking these factors into consideration, HRVs appear to be reasonably safe, with inherent characteristics that may facilitate their persistence in tumors where they can establish a productive infection. While effective OVs need to induce a strong immune response, it may be equally important to give the virus ample time to replicate and spread to enhance the anti-tumor effect. HRVs appear able to meet these criteria through their ability to induce adaptive immunity, evade neutralization through use of multiple serotypes, and avoid induction of long-term memory responses.

In this study, we investigate HRVs and establish their efficacy as a novel class of oncolytic agent. We have found that HRVs, in a wide variety of cancer cell types, are able to kill, replicate to produce a productive infection, and stimulate pro-inflammatory immune responses. Finally, we show that the viral infection promotes oncolysis and adaptive immunity, leading to regression in murine tumor models. We propose that HRVs represent a class of natural oncolytic viruses that are reasonably safe, lethal to tumor cells, conducive to anti-tumor immunity, and should be considered for further development in anticipation of clinical testing.

## Materials and Methods

### Cell culture and cell lines

Normal human epithelial melanocytes (NHEM) were purchased from Invitrogen. We have previously described the following human melanoma cell lines: A375, C32, C8161R, CACL, LOX IMVI, M14-MEL, MALME-3M, SK-MEL-28, SK-MEL-5, SK-MEL-2, SK-MEL-103, SK-MEL-147 UACC-62, UACC-257 [26]. CHL-1 and HeLa-H1 cells were purchased from American Type Culture Collection (ATCC, Manassas, VA,USA). Yale University Mouse Melanoma Exposed to Radiaiton 1.7 (YUMMER 1.7) cells were purchased from Millipore Sigma (Burlington, MA, USA) [27]. Human melanoma cell lines were cultured in RPMI 1640 supplemented with 7% fetal bovine serum (FBS) and gentamicin. YUMMER 1.7 cells were cultured in DMEM/F12 supplemented with 10% FBS, non-essential amino acids (NEAA), and gentamicin. NHEM cells were cultures in 254 media supplemented with human melanoma growth supplement, 10% FBS, NEAA, and gentamicin. Cells were cultured in humidified incubators at 37°C with 5% CO_2_.

### Western blotting

Cell lysates were prepared by washing cells once in phosphate buffered saline (PBS) followed by the addition of 0.5ml of 0.25% Trypsin to remove cells from the tissue culture plates. Cells were collected in RPMI media (7% FBS; gentamicin) washed in PBS, and re-suspended in cold RIPA buffer with protease inhibitors and EDTA for protein extraction. Cell were lysed by agitation on a rotating platform at 4°C for 20 minutes and clarified at 17,000xg for 10 minutes at 4°C. Protein was quantified using a bicinchoninic acid (BCA) protein assay (Pierce; 23225) according to manufacturer’s instructions. Following quantification, 10µg of protein were diluted in 4x lithium dodecyl sulfate (LDS) buffer (Life Technologies; NP0008) with dithiothreitol, denatured at 95°C for 5 minutes, and loaded on to 4-12% Bis-Tris gels (Life Technologies; NP0321BOX/NP0323BOX). Gels were run at 200 volts for 40 minutes. Proteins were transferred in NuPAGE transfer buffer (Thermo Fisher; NP0006) to a nitrocellulose membrane (Bio-Rad; 162-0232) at 90 volts for 90 minutes and blocked in 5% non-fat dry milk in 0.05% TBS-T for 30 minutes at RT. Membranes were washed several times in 0.05% TBS-T, and probed with primary antibodies diluted in 5% BSA with agitation at 4°C. The membrane was washed several times in 0.05% TBS-T. Anti-mouse (Cell Signaling Technologies; 7076S) and anti-rabbit (Cell Signaling Technologies; 7074S) secondary antibodies were diluted 1:1000 in 0.05% TBS-T and used to probe the membrane for 1-2 hours with agitation at 4°C. The membrane was then washed several times in 0.05% TBS-T and proteins were detected using enhanced chemiluminescence (ECL) reagent (GE Healthcare; RPN2106). Antibodies used include ICAM-1 (Cell Signaling Technologies; Rabbit; 4915), HA (Cell Signaling Technologies; Rabbit; 3274), and GAPDH (Millipore; Mouse; MAB374).

### Reverse-transcription polymerase chain reaction (RT-PCR)

RT-PCR to detect HRV was carried out in two steps. Reverse transcription was performed on RNA samples using the ProtoScript II first strand cDNA synthesis kit (New England BioLabs; E6560S) with HRV serotype-specific reverse primers (Rev 5’- -3’). The reaction was carried out at 25°C for 5 minutes, then 42°C for 1 hour followed by inactivation of the enzyme at 80°C for 5 minutes. Once cDNAs were generated for each sample, HRV amplicons (408bp) were generated by polymerase chain reactions (PCR) using EconoTaq PLUS Green 2X Master Mix (Lucigen; 30033-1) and HRV serotype-specific primers (Fwd 5’- -3’; Rev 5’- -3’). The PCR was run with an initial 95°C denaturing step for 10 minutes, followed by 21 cycles of denaturing at 95°C for 20 seconds, annealing at 55°C for 20 seconds, and elongation at 68°C for 30 seconds. The PCR was concluded with a final elongation step of 72°C for 5 minutes prior to cool down to 4°C. Amplified DNA was detected by running 25µl of the PCR product on a 1% agarose gel stained with ethidium bromide and imaged with a UV illuminator.

### HRV propagation, purification, and titration

HRVs were propagated in HeLa-H1 cells (ATCC). A 10cm dish of HeLa-H1 cells was infected at 90-100% confluence with 1×10^6^ IFU (MOI 0.1) for 8-24 hours, whereupon cells were observed for signs of cytopathological effect (CPE). Upon confirmation of <50% CPE, cells were scraped into the culture media, collected into conical tubes, and flash frozen using a methanol-dry ice bath. Virus was released from infected cells by thawing in a 47°C water bath followed by rigorous vortexing for 30 seconds. Freeze-thaw cycles were carried out three times on samples before being cleared at 5,000rpm for 10 minutes at 4°C. Viral supernatants were then 0.8µm filtered into clean tubes. Viral stocks were stored at -80C in aliquots.

To generate purified viral stocks, HeLa-H1 cells in suspension culture were allowed to grow between 5×10^5^ and 1.5×10^6^ cells/mL in 800mL of S-MEM media [10% FBS, Non-essential amino acids (NEAA), Pleuronic-F68, L-glutamine, gentamicin]. Cells were then pelleted at 500xg for 10 minutes, re-suspended in 45mL of media, and infected with 35mL of propagated HRV stock. The infection was carried out at 37°C for 1 hour with shaking at 70rpm. 80mL of fresh S-MEM was then added to the flask and the infection proceeded at 37°C with shaking at 120rpm until limited (<50%) CPE was observed. The cell suspension was freeze-thawed and cleared as previously described and aliquoted equally into 25×89mm SW 28 ultracentrifuge tubes (Beckman Coulter; 326823). 1.5mL of a buffered 30% sucrose solution (20mM Tris, 1M NaCl) was added to the bottom of each tube to create a sucrose cushion. Tubes were weighed in rotor buckets and balanced with PBS. HRV cultures were ultracentrifuged at 25,000rpm for 4 hours at 4°C. Supernatant was decanted and the virus pellets were suspended in 100µl of PBS, covered, and placed at 4°C overnight to allow the pellets to dissolve. The pellets were then combined, suspended in 4mL of PBS, and stored at -80°C in 50µl aliquots.

To determine viral titer, HeLa-H1 cells were seeded in two 6-well cell culture dishes at 50-60% confluence. A ten-fold serial dilution of the viral stock was prepared in Opti-MEM media and cells were infected with 500µl of each dilution for 2 hours at 37°C. Following the infection, cells were washed in PBS to remove unincorporated virus and then incubated in 1mL of normal culture media (RPMI; 7% FBS, gentamicin) for 72 hours at 37°C. RNA was then purified from the cells by TRIzol (ThermoFisher Scientific; <u>15596018</u>) extraction according to manufacturer’s instructions. Cells were washed in PBS and 400µl of TRIzol reagent was added to each well. Wells were rinsed vigorously with the TRIzol and cells were collected into tubes and incubated at RT for 5 minutes. 80µl of chloroform was then added and samples incubated for an additional 3 minutes. Separation of the aqueous, inorganic, and organic layers was carried out by centrifugation at 12,000xg for 15 minutes at 4°C. The aqueous phase was then transferred to a tube containing 15µg of GlycoBlue (ThermoFisher Scientific; AM9515). RNA was precipitated with 200µl of isopropanol and incubated for 10 minutes at room temperature prior to being cleared at 12,000xg for 10 minutes at 4°C. The supernatant was discarded and the pellet was re-suspended in 75% ethanol and vortexed prior to centrifugation at 7,500xg for 5 minutes at 4°C. The supernatant was removed and the pellet was allowed to air-dry at room temperature for 10 minutes. Samples were re-suspended in 20µl of RNase-free water and heated at 55°C for 15 minutes. RT-PCR was then performed to detect HRVs in the extracted RNA. RNAs were stored at -80°C.

### Flow cytometry

Cell death assays were performed using the Attune Nxt Acoustic Focusing Cytometer (Life Technologies). Cells were seeded in 6-well plates at a density of 1×10^6^ cells per well. The following day cells were infected with HRVs at an MOI of 0, 0.1, 1.0, or 10 in phenol red-free RPMI (7% FBS; gentamicin) for 1 hour. Cells were then washed with PBS, re-fed fresh media, and incubated for 24 to 48 hours. At the experimental endpoint media was collected and combined with a PBS wash for each sample. Following treatment with 0.25% trypsin, adherent cells were re-suspended in the media/PBS mixture from each well. Cells were pelleted by centrifugation at 500xg for 10 minutes at 4°C and re-suspended in 1ml of 0.2µM SYTOX Green (Invitrogen; S7020) reagent diluted 1:1000 in PBS from a 5mM stock. Cells were stained in the dark at room temperature (RT) for 20 minutes prior to being stored on ice and run on the Attune. Cells were detected using a blue laser (488nm excitation BL1 530/30 bandpass filter) and analyzed for uptake of the dye indicative of cell death. Cell death was quantitated by averaging the percentage of cell death in three replicates. Error bars were generated based on the calculated standard deviation of the replicates.

Tumor profiling was performed using the BD LSRFortessa flow cytometer. Tumor initiation treatments were performed as described. At the experimental endpoint, mice were euthanized and tumors were collected into pre-weighed tubes containing 5mL of serum-free RPMI media. Samples were then weighed again to determine the mass of each tumor collected. Tumors were dissociated in the RPMI using scissors and forceps and by grinding the tumors between frosted glass slides. 250µl of 20x Collagenase IV/DNAse was then added and dissociated tumor sample were incubated with gentle shaking at 37°C for 45 minutes. Following the collagenase digestion, samples were 40µm filtered followed by an additional wash of 5ml of serum-free RPMI into the same collection tube. Cells were collected by centrifugation at 1500rpm for 5 minutes at 4°C and decanted prior to resuspension in 1mL of ACK (Ammonium-Chloride-Potassium; pH 7.22) buffer. Cells were again cleared by centrifugation as before and resuspended in 300µl of RPMI (2% FBS) and 3 wells/mouse were plated in round-bottomed 96-well plates at 100ul/well. Cells were pelleted in the plate at 2000rpm for 1 minute and stained using three different panels, including myeloid [F480 (APC), CD45 (AF488), CD86 (BV421), Ly6C (BV510), Live/Dead Aqua (BV605), CD80 (BV650), CD11b (BV711), MHCII (IA/IE) (BV786), B220 (PE), CD11c (PE-CF594), CD8a (PECy7), Fc Block], lymphoid [CD44 (APCe780), CD45 (AF488), CD127 (PerCP Cy5.5), Live/Dead Aqua (BV510), PD-1 (BV605), CD4 (BV650), KLRG1 (BV711), CD8a (BV786), CXCR6 (PE), CD62L (PE-CF594)], and transcription Factor panels [FoxP3 (APC), BCL6 (APC Cy7), CD45 (AF488), GrzB (BV421), Live/Dead Aqua (BV510), Tbet (BV605), CD4 (BV650), CD8a (BV786), TCF1 (PE)].

Staining for myeloid and lymphoid panels was carried out on ice in the dark for 30 minutes and clarified at 2000rpm for 1 minute. Stained samples were then washed three times in 1xPBS and clarified under the same conditions. Cells were stained again with live/dead stain (1:1000 in PBS) for 10 minutes in the dark at RT. Following live/dead staining, samples were washed three times in FACS buffer and clarified as before. Myeloid and Lymphoid panels were then fixed in Cytofix/Cytoperm for 5 minutes at RT in the dark, washed three times in FACS buffer, and resuspended in 200µl of FACS buffer prior to overnight incubation at 4°C in the dark.

Cells used for the transcription factor panel were permeabilized in FOXP3 perm buffer (eBiosciences) and washed in FOXP3 perm wash (eBiosciences). They were fixed in FOXP3 Fix buffer for 10 minutes at RT in the dark followed by additional washes in FOXP3 perm buffer. Antibodies for the transcription factor panel were suspended in FOXP3 perm buffer and cells were stained on ice for 30 minutes. Cells were then washed once in perm buffer and twice in FACS buffer, followed by a final resuspension in FACS buffer and incubated at 4°C overnight in the dark.

Data was analyzed using a two-way ANOVA test with a Tukey post-test. *P* values <.05 were considered significant (*P* <.05*, <.01**, <.001***,<.0001****).

### HRV enzyme linked immunosorbent assay (ELISA)

A 96-well ELISA plate (Costar; 3590) was coated with HRV by incubating each well in 2×10^5^ IFU of HRV in 100µl of PBS for 1 hour at RT. Coated plates were stored overnight at 4°C. HRV was discarded and wells were incubated in 250µl of blocking buffer (ThermoFisher; TBS Buffer 28358, 0.5% Tween, 0.5% BSA) for 1 hour at RT. Blocking buffer was then discarded and mouse serum samples were diluted 1:50, 1:500, 1:5000, and 1:50000 in blocking buffer and incubated in the antigen coated wells for 1 hour at RT. The plate was then washed three times in 250µl of wash buffer (ThermoFisher; TBS Buffer 28358, 0.5% Tween) and incubated in HRP-conjugated goat anti-mouse IgG secondary antibody (ThermoFisher, 31430) diluted 1:5000 in blocking buffer. Wells were incubated in 100µl of secondary antibody for an hour at RT. Following four additional wash steps, 50µl of TMB reagent (ThermoFisher; 34028) was added to each well. After 15-30 minutes, 50µl of stop solution (ThermoFisher; SS04) was added to the plate and the absorbance at 450nm was quantified using a plate reader.

### Mice and injectables

C57BL6 (BL6) mice were obtained from the Charles River Laboratories (Raleigh, NC, USA). Congenic ICAM-1 mice were generated by crossing C.FVB-Tg(ICAM1)4Grom/J mice (The Jackson Laboratory) with C57BL6 mice to produce heterozygous ICAM-1 transgenic mice [28, 29]. The presence of the transgenic ICAM-1 allele was confirmed by PCR using human ICAM-1 specific primers (Fwd 5’-GCTGGTGAGGAGAGATCACC-3’; Rev 5’-ACCTGGGCCTCCGAGACTGG-3’). Transgenic F1 offspring were then backcrossed to C57BL6 mice. This process of selection and backcrossing was carried out for 7 generations. C57BL6 ICAM-1 mice were injected with 2×10^6^ YUMM 2.1 ICAM-1 cells in order to determine whether they were sufficiently syngeneic to allow tumor growth. High tumor penetrance (90-100%) was observed in this model. C57BL6 ICAM-1 mice were then crossed in order to make the transgene homozygous.

Tumors were initiated in 6-9 week old mice by subcutaneous flank injections of 10×10^6^ YUMMER 1.7 ICAM-1 cells. Prior to injection, cells were washed once in PBS and once in Hank’s balanced salt solution (HBSS) before a final suspension in 100µl of HBSS. Tumor re-challenge experiments were carried out by preparing and injecting cells in the same manner in the opposing flank.

HRV intratumoral injections were administered when tumors reached 100mm^3^. Mice received a dose of 2×10^8^ IFU of virus every other day for the first week on days 1, 3, 5, and 8, followed by injections once a week for the next four weeks on days 15, 22, 29, and 36. The _α_PD-1 antibody (clone RMP1-14; BE0146) and the isotype control antibody (clone 2A3; BE0089) were obtained from Bio X Cell (West Lebanon, NH, USA). _α_PD-1 and isotype control antibodies were delivered systemically by IP injection on days 1, 3, 5, and 8. Tumors were monitored daily and measured three times a week with electronic calipers until the longest diameter of the tumor reached 2cm or adverse tumor-related complications required animals to be euthanized. Tumor size was calculated using an elliptical estimation: V=(LxW^2^)/2, where tumor volume (V) is equal to the length (L) multiplied by the square of the width (W) divided by two.

Tumor responses were classified as progressive disease (PD), partial response (PR), stable disease (SD), or complete response (CR) according to RECIST 1.1 criteria [30]. According to this criteria, PD is determined by a 20% increase in the diameter of target lesion (with no CR, PR, or SD documented before increased disease). PR is classified as a 30% decrease in the longest diameter of the target lesion with recurrence, whereas SD is when neither PR nor PD criteria are met from the smallest diameter since treatment started. CR is classified as the disappearance of the target lesion with no recurrence. Survival curves were generated and analyzed using GraphPad Prism 9.0.1. Survival distributions were compared using a Log-rank (Mantel-Cox) test. *P* values <.05 were considered significant (*P* <.05*, <.01**, <.001***,<.0001****). All experiments were approved and conducted in accordance with the University of Utah Institutional Animal Care and Use Committee under protocol number 20-07006.

### Cytokine and chemokine multiplexing

Cytokine/chemokine multiplexing was performed according to the manufacturer’s instructions as described previously [31]. Mouse plasma was collected immediately following euthanasia of animal subjects. Upon sacrifice, 200µl of blood was collected into tubes coated with 10µl of 0.25mM EDTA following cardiac puncture. Blood samples were cleared at 1600xg for 10 minutes at 4°C. Plasma was then transferred to clean tubes and flash frozen in a methanol-dry ice bath. Cell culture supernatants were collected 24 hours post-infection, cleared by centrifugation, and flash frozen in a methanol-dry ice bath. All samples were stored at -80°C. Samples were run using the appropriate ProcartaPlex Multiplex Immunoassay kit (Invitrogen; <u>EPXR360-26092-901</u>; Invitrogen, EPXP420-10200-901). Samples were thawed at 4°C with agitation, vortexed, and cleared at 1000xg for 10 minutes at 4°C. 25µl of each sample was then added to a 96-well assay plate and incubated with antibody-conjugated magnetic multiplexing beads for 2 hours at RT on a microplate shaker at 500rpm. Following a wash step using a magnetic plate washer, captured analytes were then probed under similar conditions with biotin-labeled detection antibodies for 30 minutes and washed prior to treatment with Strepdavidin-PE for another 30 minutes. Following the final wash steps, the beads were re-suspended in reading buffer prior to running the plate on the Luminex MAGPIX platform. Analyte differences between samples were compared using an unpaired, two-tailed t-test with GraphPad Prism software version 9.0.1. *P* values <.05 were considered significant (*P* <.05*, <.01**, <.001***, <.0001****).

### IncuCyte live cell imaging and analysis

Live cell imaging was performed using the IncuCyte platform. Cells were seeded at a density of 1.4×10^4^ cells per well and given time to adhere to the plate for 4 hours. Following cell seeding, HRV infections were performed for 1 hour, after which cells were washed with PBS and incubated at 37°C. Two pictures of each well were taken every two hours for 72 hours to assess cell death over time. Images were collected and compiled into movies using the IncuCyteZoom2016 software.

### 2.10 qRT-PCR

## Results

### HRVs infect and rapidly kill melanoma cells

HRVs exploit cell surface receptors for infection, which include ICAM-1 for major group HRVs as shown with CVA21 [32] or LDL-R family members. To test the ability of HRVs to infect melanoma, a panel of cell lines representing BRAF and NRAS-driven variants was infected with HRVA2 and HRVA45 at MOI of 0 and 1.0 for one hour, after which unincorporated virus was washed out and cells were incubated in fresh media at 37°C for 24 hours. Following another wash, RNA was extracted from cells using TRIzol and the presence of internalized HRV genomic RNAs were assayed by RT-PCR (Figure 1). Amplicons were not observed in uninfected (MOI 0) cells. Serotype specific amplicons for HRVA2 were detected in each of the other infected cell lines (MOI 1.0), including Normal Human Epithelial Melanocytes (NHEM). Major group HRVA45 was likewise able to infect all cell lines with the exception of YUMMER 1.7/YUMM 2.1 cells, which lack it’s cognate receptor. This demonstrates the ability of HRVs to be internalized by melanoma cells.

**Figure 1.**
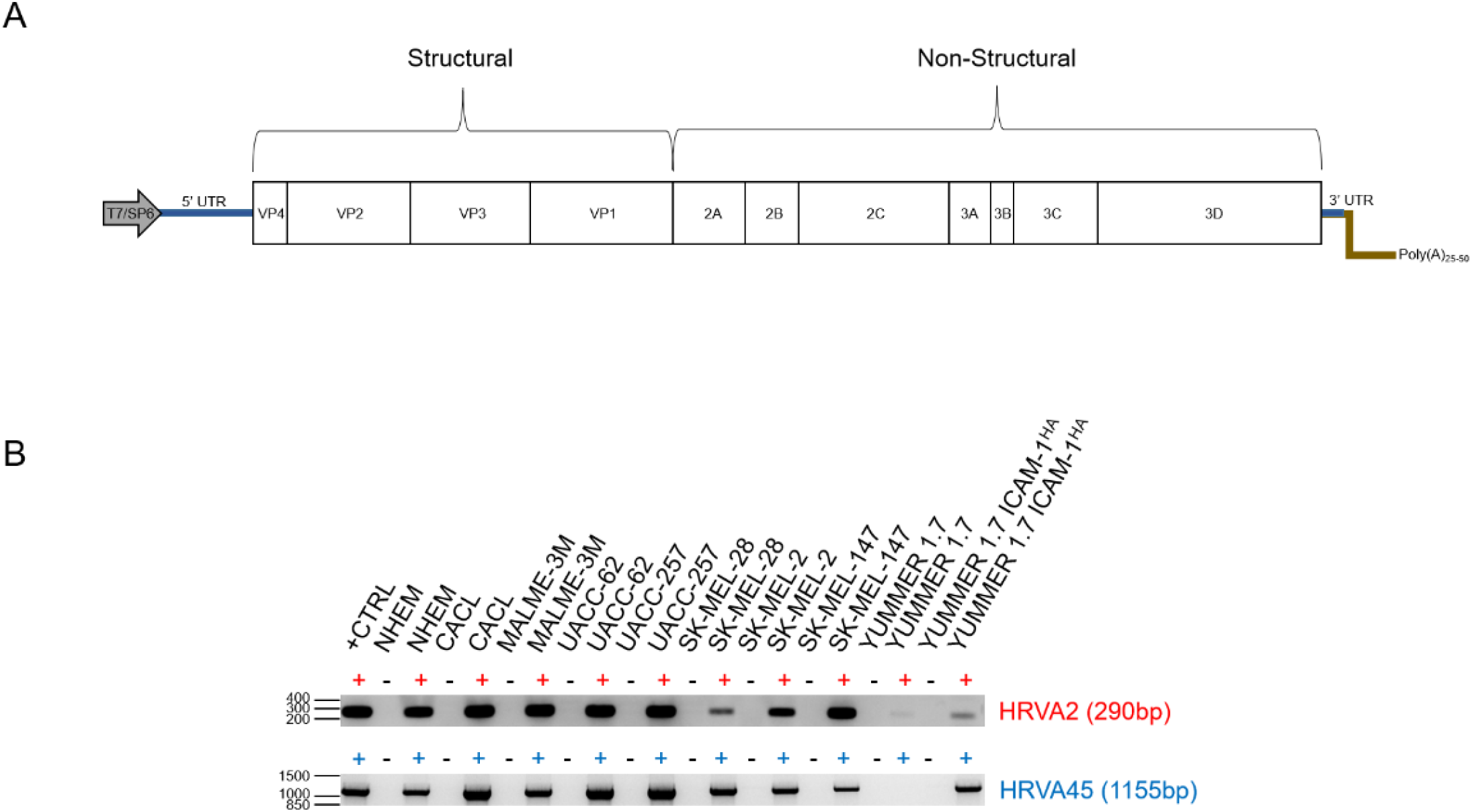
HRVs infect melanoma cells that express their cognate receptors. (A) Genomic structure of human rhinovirus. The genome is a positive-sense, single stranded RNA. Viral capsid genes VP1-VP4 are located on the 5’ end, while the non-structural components (proteases, polymerases, etc.) are on the 3’ end. (B) RT-PCR of RNA extracted from a panel of melanoma cell lines infected with HRVA2 and HRVA45. Uninfected parental lines are shown for comparison. Purified HRV stock virus was used as the positive control. The RT-PCR produces an amplicon from the 3’ end of the viral genome in the 3D^pol^.

YUMM 2.1 and YUMMER 1.7 ICAM-1/YUMM 2.1 ICAM-1 cells, which had been shown to be resistant to infection (Figure 1B). HRVA45 was able to induce cell death in YUMMER 1.7 ICAM-1 cells, but not in parental YUMMER 1.7 cells. Most susceptible cell lines exhibited high levels of cell death within 24-hours, even at low MOI (Figure 2; Video S1). While NHEM cells were sensitive to HRV-mediated cell death, they exhibited some resistance to infection relative to melanoma lines. This was further highlighted by their durability when infections were allowed to go for longer periods (Figure 2; Video S1). Thus, when HRVs are incorporated into melanoma cells they induce cell death, demonstrating their oncolytic capability.

**Figure 2.**
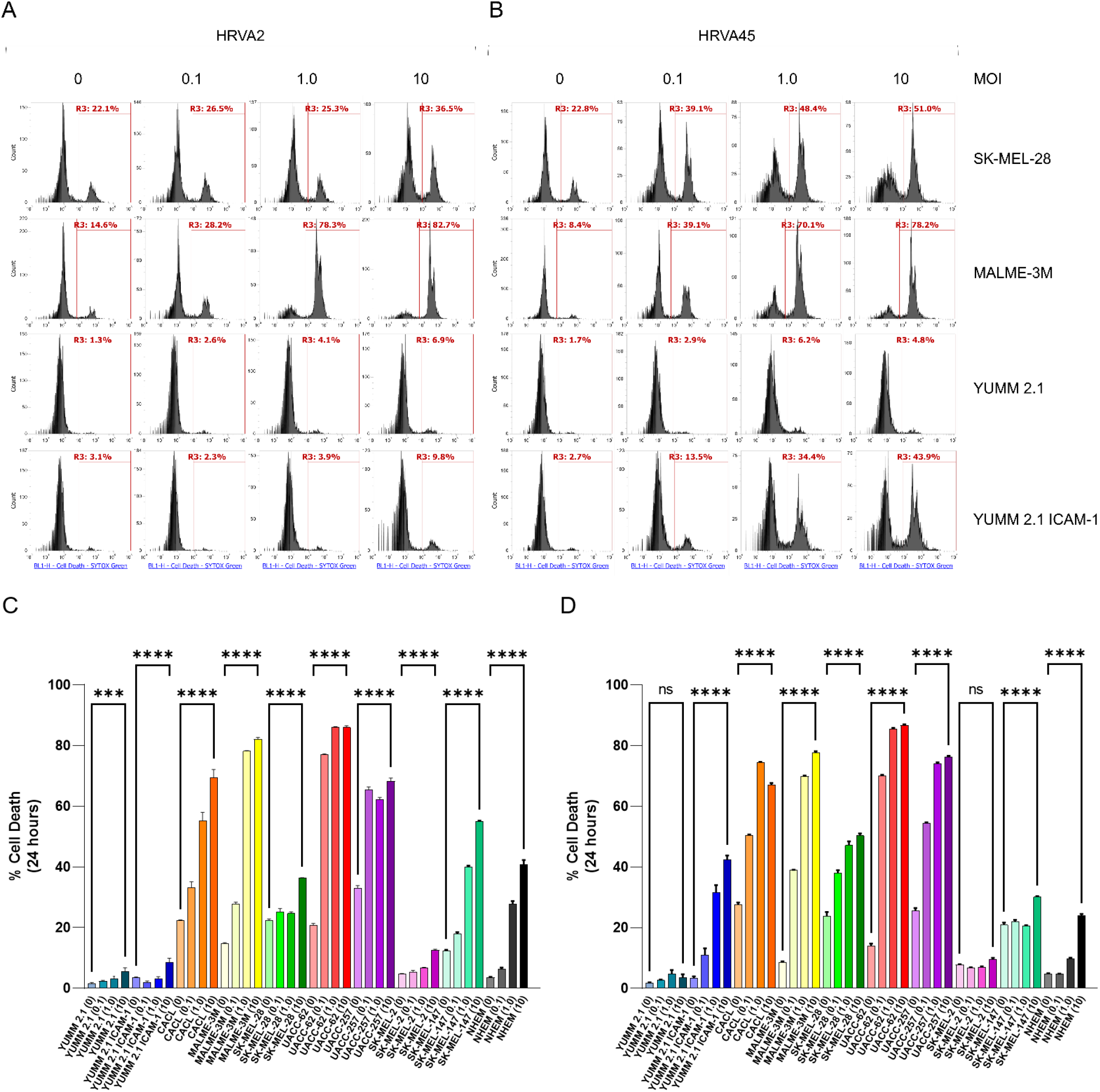
HRVs induce melanoma cell death upon infection. (A,B) Assessment of cell death following HRV infection at MOI 0, 0.1, 1.0, and 10 at 24 h by flow cytometry in representative cell lines. SYTOX Green was used to stain dead cells. (C,D) Quantitation of cell death from the flow cytometry assay represented in panel. Samples were run in triplicate and analyzed using a one-way ANOVA test for statistical differences. Error bars represent the standard deviation between replicates. p values < 0.05 were considered significant (p < 0.05 *, < 0.0001 ****).

qRT-PCR was used to determine how cell death corresponded to viral replication within infected melanoma cells. Infections with HRVA2 and HRVA45 were carried out at a MOI of 1.0 for 0, 8, and 16 hours. For each time point, cell samples and supernatants were collected to assess intracellular and extracellular virus, respectively. Viral RNA was purified by TRIzol extraction as before. Following extraction, samples were analyzed by qRT-PCR.

### HRV infection induces an inflammatory cytokine response and production of HRV-specific antibodies

As infiltration of immune cells is critical for anti-tumor immune responses, HRV’s were evaluated for their ability to induce an inflammatory response in infected cells. Melanoma cell lines and NHEMs were infected with HRVA2 or HRVA45 at an MOI of 0, 0.1, and 1.0 for 24 hours, after which cell-free supernatants were collected and probed using a cytokine multiplexing approach. Cytokine levels tended to increase upon infection of NHEMs, resulting in significant increases in [cytokines], though the magnitude of those increases was less than in melanoma cells (Figure 4A). There was a significant increase in [cytokines] in [cell lines] upon infection with either HRV, and many additional cytokines that did not attain statistical significance were nevertheless higher in infected cells (Figure 4A). Cytokine levels were higher in cells infected with a lower MOI. This is evidence that, whether in melanocytes or tumor cells, HRV infection drives a pro-inflammatory response.

The ability of HRVs to induce an adaptive immune response in a murine host were also assessed. Serum collected from mice inoculated with HRVA2 or HRVA45 was tested against serum of naïve mice for neutralizing antibodies using an ELISA assay (Figure 2B). A 96-well plate was coated with each HRV antigen and incubated with serum samples diluted 1:50, 1:500, 1:5000, and 1:50000. Following antibody capture, wells were washed and treated with anti-mouse IgG detection antibody conjugated to horseradish peroxidase (HRP). Absorbance was quantified using a plate reader. The detection antibody yielded no signal in the absence of serum antibodies, and very little signal was detected in wells treated with the serum of naïve mice. Conversely, wells treated with serum from mice inoculated with either HRV exhibited strong absorbance, characteristic of the binding of virus-specific antibodies. Although this signal was somewhat diminished two weeks later, the presence of HRV antibodies was readily detectable. These antibodies are indicative of an adaptive immune response to HRV infection.

### HRV exposure is well tolerated in a murine melanoma model that is susceptible to HRV infection of normal mouse tissues

The safety of the virus was then evaluated *in vivo*. As HRVs are human pathogens and mice are refractory to infection, a murine model was developed to more accurately recapitulate human susceptibility to viral infection (Figure 3A). C57BL6 mice were crossed with mice carrying a transgenic ICAM-1 gene under the control of endogenous regulatory elements [28, 29]. Transgenic ICAM-1 mice were backcrossed onto the C57BL6 background until they were sufficiently syngeneic to support growth of YUMMER 1.7 ICAM-1 tumors with high penetrance, after which mice were made homozygous for the transgene. In this model, the endogenous expression of ICAM-1 closely mirrors the regulation of ICAM-1 in human tissues, where it is highly prevalent in the lung and spleen and minimally expressed in other tissues such as the kidney, heart, and brain (Figure 3B) [28, 29]. Expression of ICAM-1 renders normal murine tissue vulnerable to HRV infection, allowing murine hosts to be used as a surrogate for HRV induced pathogenesis.

**Figure 3.**
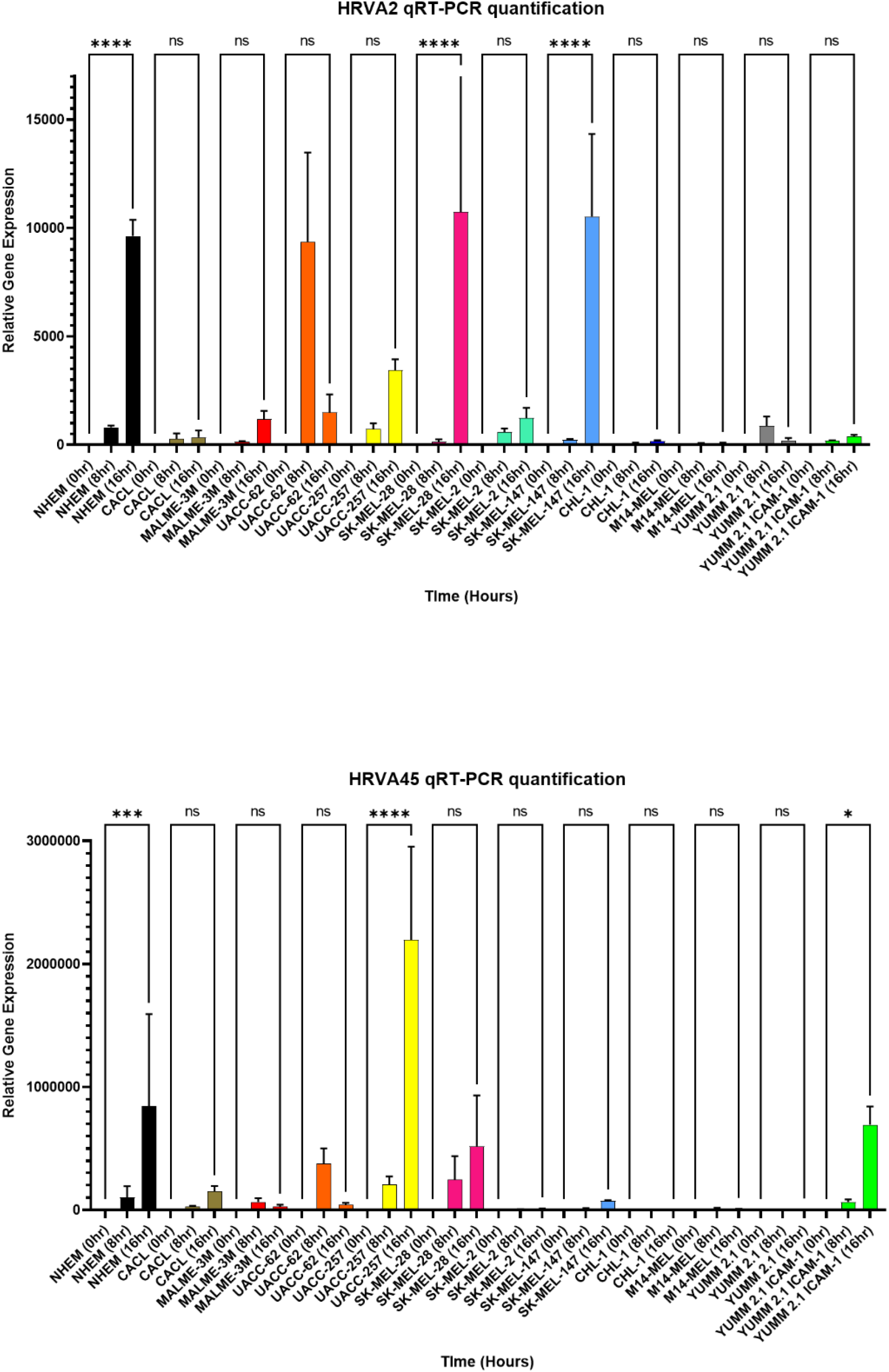
HRVs exhibit variable replicative efficiency in melanoma cells. The presence of intracellular or extracellular HRV was quantified using qRT-PCR. Relative gene expression was determined by normalizing infected cells to uninfected cells for each virus.

**Figure 4.**
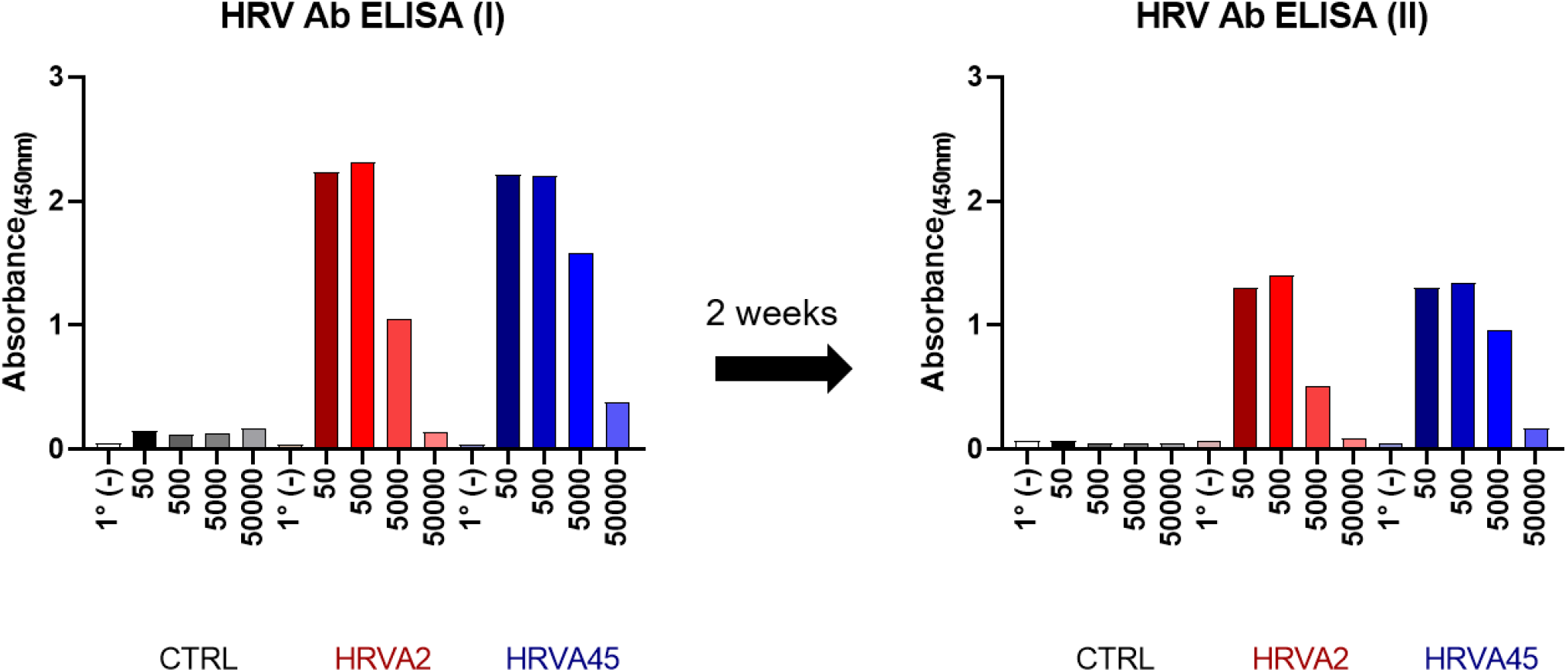
HRVs induce inflammatory signaling and adaptive immune responses. (A) Cytokine multiplexing in melanoma cell lines. Supernatant samples collected from cells infected with HRVA2 or HRVA45 at MOI 0, 0.1, or 1.0 for 24 hours. *P* values <.05 were considered significant (*P* <.05*, <.01**, <.001***, <.0001****). (B) RT-PCR time course of HRVA45 delivered systemically IP and extracted from the blood of inoculated mice. (C) ELISA test for HRVA2 and HRVA45 antibodies in the serum of naïve and inoculated mice immediately and 2 weeks post-inoculation.

**Figure 5.**
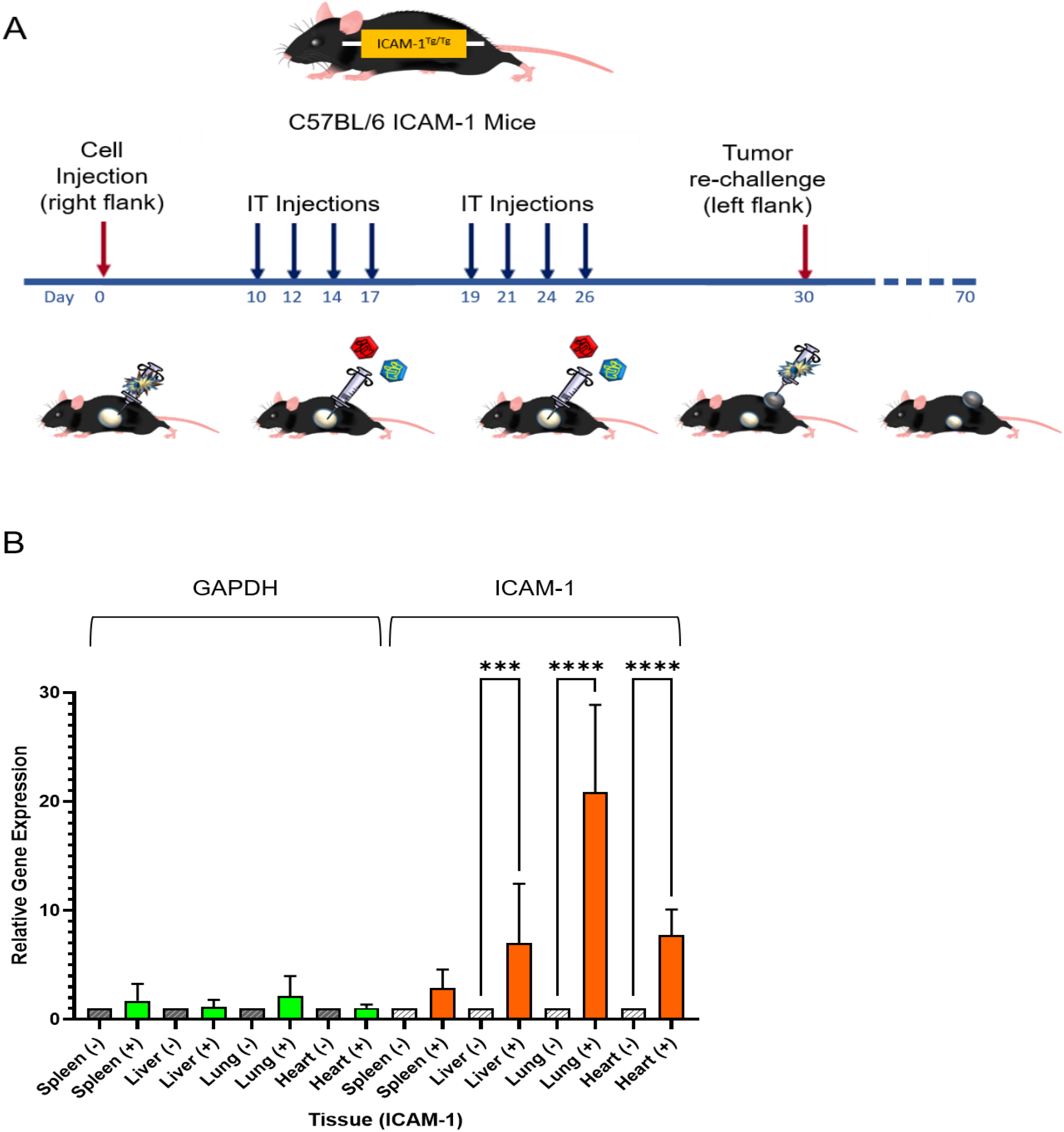
HRVs induce inflammatory signaling and adaptive immune responses. (A) Cytokine multiplexing in melanoma cell lines. Supernatant samples collected from cells infected with HRVA2 or HRVA45 at MOI 0, 0.1, or 1.0 for 24 hours. *P* values <.05 were considered significant (*P* <.05*, <.01**, <.001***, <.0001****). (B) RT-PCR time course of HRVA45 delivered systemically IP and extracted from the blood of inoculated mice. (C) ELISA test for HRVA2 and HRVA45 antibodies in the serum of naïve and inoculated mice immediately and 2 weeks post-inoculation.

**Figure 6.**
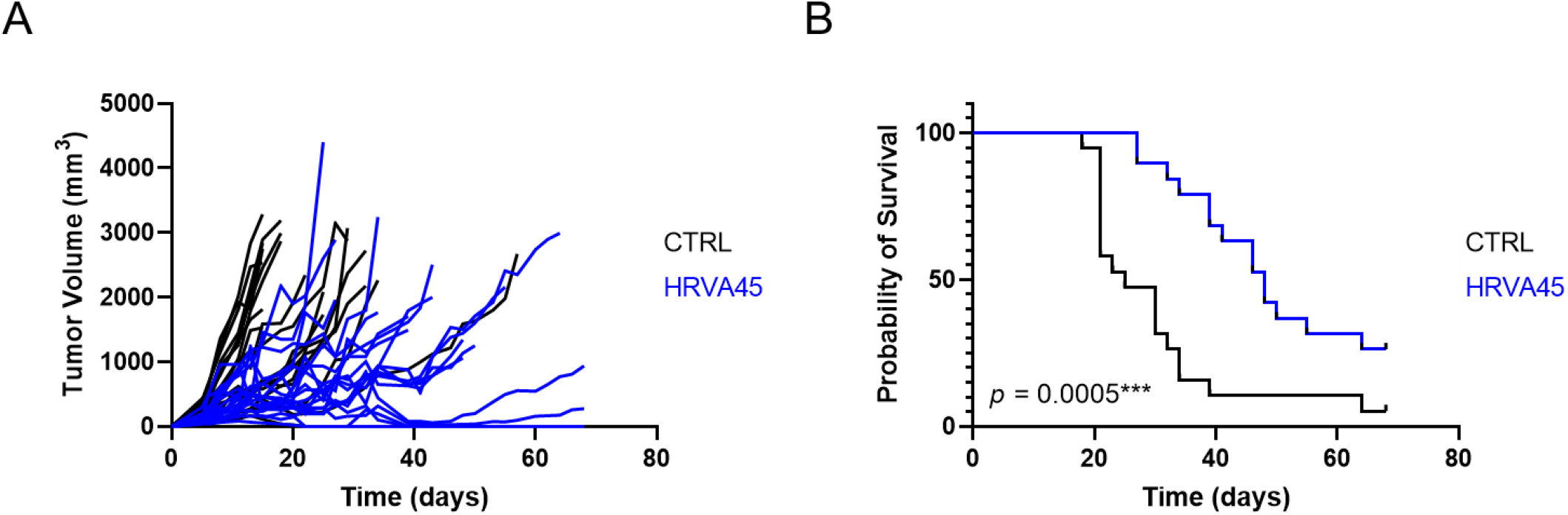
Tumor responses and overall survival in C57BL/6 ICAM-1 mice treated with IT saline (CTRL) or HRVA45. (**A**) Tumor growth measured in CTRL (n = 19) and HRVA45 (n=19) treated mice over 70 days. Responses are classified as progressive disease (PD), partial response (PR), or complete response (CR) according to RECIST 1.1 criteria. (**B**) Survival of CTRL (n = 19) and HRVA45 (n=19) treated mice over 70 days. Survival distributions were compared using a Log-rank (Mantel-Cox) test. p values < 0.05 were considered significant (p < 0.05 *, < 0.001 ***, < 0.0001 ****).

The health of C57BL6 ICAM-1 mice was evaluated following systemic delivery of 2×10^7^ IFU of HRVA2 or HRVA45 by intraperitoneal (IP) injection. None of the mice exhibited deleterious symptoms, even after repeated dosing in this manner. Similarly, when YUMMER 1.7 ICAM-1 tumors were initiated, repeated dosing of these tumors with intratumoral injection of HRV had no visible effect on mouse health.

To determine how quickly HRVs are cleared from the animals, blood was drawn from naïve mice and those inoculated with 2×10^7^ IFU of HRV by IP injection at 1, 2, 4, and 8 hours post-injection. HRV was detected in the blood using RT-PCR as described previously, where RNA levels were first normalized prior to reverse transcription. HRVs were detected in the blood out to 8 hours following systemic delivery, with viral load decreasing at that time point (Figure 3C).

Taken together, this data demonstrates that C57BL6 ICAM-1 mice are susceptible to HRV infection but are refractory to HRV-induced pathogenesis. Whether by localized IT injection or systemic exposure to the virus, mouse subjects remain healthy and are capable of clearing the virus relatively rapidly. This safety profile supports further investigation of HRVs for their efficacy as oncolytic agents.

### HRVs do not drive tumor regression in an immuno-deficient murine model of melanoma

In order to assess the oncolytic activity of HRV independent of the immune system, tumors were generated in immune-compromised nude mice by injecting 2×10^6^ SK-MEL-28 cells subcutaneously in the right flank. In this experiment, HRV was tested against CVA21, a known oncolytic picornavirus that has demonstrated efficacy in murine tumor models and clinical benefit (CVA21 paper reference, VLA-007 CALM). We have shown that SK-MEL-28 cells are sensitive to infection with both viruses. When tumors reached 50mm^3^ they were treated with 2×10^8^ IFU of either CVA21 or HRVA2 on days 1, 3, 5, 8 in the first week and once a week for the next four weeks on days 15, 22, 29, and 36.

Mice were monitored daily and tumor growth was measured 3 times per week. SK-MEL-28 tumors grew steadily in control mice, with all animals requiring sacrifice due to tumor burden by 40 days (Figure 4). Similarly, tumors treated with either HRV or CVA21 exhibited little response to viral infection, with mice largely needing to be euthanized within 40 days, and all mice requiring sacrifice due to tumor burden.

### 3.5 HRVs drive tumor regression in an immuno-competent murine model of melanoma and responses are enhanced by using HRVs in combination and with _α_PD-1

As HRVs demonstrated robust oncolytic activity *in vitro*, the anti-tumor activity of HRV was then examined in an immuno-competent murine melanoma model. C57BL6 ICAM-1 mice were injected subcutaneously with 1×10^7^ YUMMER 1.7 ICAM-1 cells in the right flank. When tumors reached 100mm^3^, mice were given a control saline injection, or 2×10^7^ IFU of CVA21 or HRVA45 via IT injection following the dosing regimen previously described. The experimental endpoint for the study was 70 days.

---

**Figure S1.**
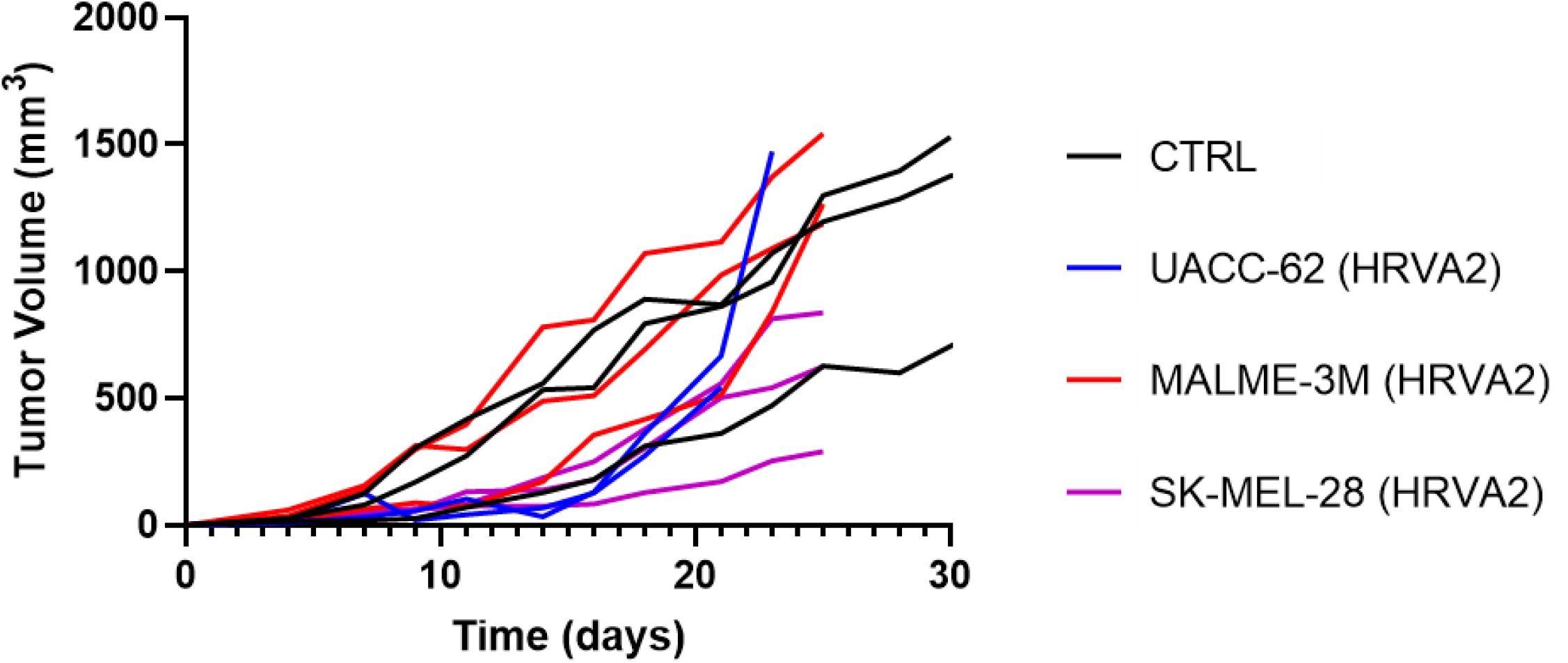
Tumor responses in nude mice treated with HRV.

